# NLP-based tools for localization of the Epileptogenic Zone in patients with drug-resistant focal epilepsy

**DOI:** 10.1101/2022.11.18.516120

**Authors:** Sara Mora, Rosanna Turrisi, Lorenzo Chiarella, Laura Tassi, Roberto Mai, Lino Nobili, Annalisa Barla, Gabriele Arnulfo

## Abstract

**Background:** Drug-resistant focal epilepsy, defined by failure of two antiepileptic drugs, affects about 30% of patients with epilepsy. Epilepsy surgery may represent an alternative options for this population. However, defining the epileptogenic zone to be surgically removed requires highly specialised medical expertise as well as advanced technologies. The aim of this work is building a cost-effective support system based on text, in particular based on the semiological descriptions of the seizures (temporal vs extratemporal lobe; right vs left hemisphere), in order to predict the localization of seizure origin.

**Methods:** Among a population of 121 surgically treated and seizure-free drug-resistant patients suffering with focal epilepsy, recruited at the Niguarda Hospital in Milan, we extracted a total number of 509 descriptions of seizures. After a data pre-processing phase, we used natural language processing tools to build numerical representations of the seizures descriptions, both using embedding and count-based methods. We then used machine learning models performing a binary classification into right/left and temporal/extra-temporal.

**Results:** All predictive models show a better performance when using the representations relying on embedding models respect to count-based ones. Between all the combinations of representations and classifiers, the best performance obtained in terms of F1-score is 84.7% ***±*** 0.6.

**Discussion:** This preliminary work reached encouraging results considering both localization tasks. The main advantage is that no specific knowledge about epilepsy is used to build the models, rendering our pipeline applicable also in other scenarios. The major limitation lies in the fact that the text is highly specific to the writer.

## 1 Background

### 1.1 Problem statement

Epilepsy is one of the most common neurological disorders. According to the *World Health Organization* (WHO) [1], epilepsy affects almost 50 million individuals worldwide and up to 10% of people have one seizure during their lifetime. In the daily clinical practice, epileptologists need to consider multiple different aspects and symptoms, in order to first determine the syndromic epilepsy diagnosis, and then define the procedures according to the specific epilepsy type. While only very few types of epilepsy can afford a slow pace of action, the majority of them need instead to be rapidly processed so that seizures may be controlled providing patients with a better quality of life. However, diagnosis and classification of epilepsy is not straightforward. People suffering from epilepsy may experience very different types of seizures, with a main distinction between focal and generalized onset. Although the complex process of epilepsy diagnosis should lead to the prescription of the most appropriate pharmacological treatment, there is a group (more than one third) of patients that do not respond to *Anti-Seizure Medications* (ASMs). In particular, if they do not respond to at least 2 appropriate ASMs, their epilepsy is defined as *“drug-resistant epilepsy”* (DRE) according to the *International League Against Epilepsy* (ILAE)^1^. In focal DRE, when a clear cut demarcated *epileptogenic zone* (EZ) can be identified, brain surgery must be considered. However, despite the evidence suggesting the favourable implication of early surgery, it generally takes several years before a patient suffering from DRE is addressed to surgery evaluation [2–4]. This issue is even more relevant in DRE children, who may benefit from epilepsy surgery at an early age and significantly improve their future well-being. Indeed, specific studies on children report delays of about 5 years between epilepsy onset and surgery [5]. Further, in the event that a surgical option is considered advisable, the patient has to go through a complex diagnostic-therapeutic process. This includes a patient-specific work-flow, including in 1/3 of the patients even invasive investigations, to accurately identify the EZ and to establish if it can be removed without causing new neurological and/or cognitive deficits. Pre-surgical examinations include, among others, *video-Electro-Encephalogram* (video EEG), *Magnetic Resonance Imaging* (MRI), *Positron Emission Tomography* (PET), and in more complicated cases, invasive recordings. In particular, the video-EEG is an important tool allowing the observation of crucial clinical signs, useful for localizing the onset of seizures and correctly driving the workflow of the pre-surgical evaluation. The clinical interpretation of the video recorded seizures requires specific knowledge which is obtained after a long period of training.

All these diagnostic procedures are often very expensive, requiring highly specialized personnel and they may not be available in all centres.

Therefore, an advanced system able to suggest an accurate probabilistic seizure origin localization based on non-invasive, cost-effective, and widely available information in primary-care units could diminish the delay in the beginning of the surgical process. This would also significantly impact patient quality of life and healthcare expenditures.

### 1.2 State of the art

As in all scientific domains, clinical and medical knowledge evolves over time. Thanks to the rapid adoption of information technologies in the clinical settings, the amount of patient-related information available in electronic form is growing incredibly fast. Enabling timely access to this clinical information is extremely important for several reasons, above all to improve patient care and to facilitate knowledge discovery. *Electronic Medical Records* (EMRs) have progressively become crucial components of clinical practice and public health. Medical centres daily produce a vast amount of textual reports which are seldom used as a source of information for investigation in medical research, as their management and processing demands a huge amount of human effort and time.

During the last years the problem of dealing with unstructured data became massive and involved several research studies not only in biomedical contexts [6, 7] but also in many different fields such as sentiment analysis [8], information extraction from textual data [9] and speech synthesis [10].

*Natural Language Processing* (NLP) is a branch of artificial intelligence, computer science and linguistics developed with the aim of transforming texts written in natural language into annotated knowledge that computers can process and analyze. This enabled the use of statistics and *Machine Learning* (ML)-based methods also on unstructured data. The use of NLP for extracting information from EMRs is a growing area of interest throughout medicine, resulting in several NLP-based pipelines that have been already tailored and applied in various use cases, especially in primary care units. For example [11] applied processes of pattern recognition to identify key signs of *Kawasaki disease* (KD) while [12] used the analysis of keywords’ frequency to link the most important clinical concepts to the diagnosis of rare diseases. Moreover, NLP tools in synergy with established ML methods, such as *Support Vector Machines* (SVM) [13, 14] and *Logistic Regression* [15, 16], often support the process of differential diagnosis [17, 18]. As regards the epilepsy field, literature reports different tools developed as answers to specific needs. For instance, the authors of [19] present a decision support system that exploits 7 questions addressed to the patients providing an accurate classification of seizure types. Epifinder [20] is a clinical decision support system for epilepsy diagnosis which, given some keywords extracted from the semiological descriptions of the seizures, returns the probability to being affected by epilepsy. ExECT (*“Extraction of Epilepsy Clinical Text”*) is an information extraction tool, which fills the routinely collected data with missing information such as epilepsy type, seizure frequency and neurological investigations [21].

### 1.3 Our contribution

As time plays a key role in the EZ localization process in DRE patients, the aim of this work is building a cost-effective support system based on text, that is the simplest and always available type of information. Therefore we aim at transforming the EMR from a mere collection of clinical information into a powerful instrument for clinical investigation. The developed system takes as input the raw text containing the description of the seizures and it predicts the probability of the EZ to be in the temporal/extra-temporal lobes and left/right hemispheres. One of the main challenges that we faced during the development of this pipeline is that not only most of the NLP-based tools are specifically built for the English language but also no Italian ontology on epilepsy is available. An ontology is an “explicit specification of a conceptualisation” [22] and it provides links between concepts and relations that are precious in an information processing task [23]. In particular, an ontology can be useful in the text normalization process, first because it is a collection of concepts and words strictly linked to the specific context and then it also provides properties of these concepts, for example known synonyms. To the best of our knowledge, this is the first work that uses the semiological descriptions of seizures written in Italian language to identify the EZ.

The rest of this paper is organized as follows. In section “Methods” we describe the dataset that we used for the analysis, the NLP and ML-based steps that compose the developed pipeline, and the three representations of the textual dataset that we obtained. In section “Results” we present the performance of the three representations on the two tasks (temporal/extra-temporal and left/right) cited above with ten-fold cross-validation on the learning set. We also performed classification on a blind test set in order to test the ability in generalization of the models. In section “Discussion” we illustrate the advantages and the limitations of our approach. Finally, we summarize our results in the “Conclusion” section.

## 2 Methods

The purpose of this study is to predict the localization (temporal/extratemporal and right/left) of the seizure origin in DRE patients based on the semiological descriptions of the seizures. Fig. 1 shows the sequence of steps we performed. First, we had to pre-process the collected textual data to anonymize, standardize and filter it *(Sec. Data preparation)*. Then, we extracted three possible numerical representations of the seizures descriptions based on Bag of Words and Word Embedding methods *(Sec. NLP & Numerical Representation)*. Finally, we applied machine learning models, using the numerical representations as inputs to predict in a first step the brain region and then the lateralization of the EZ *(Sec. ML methods for classification)*.

**Fig. 1.**
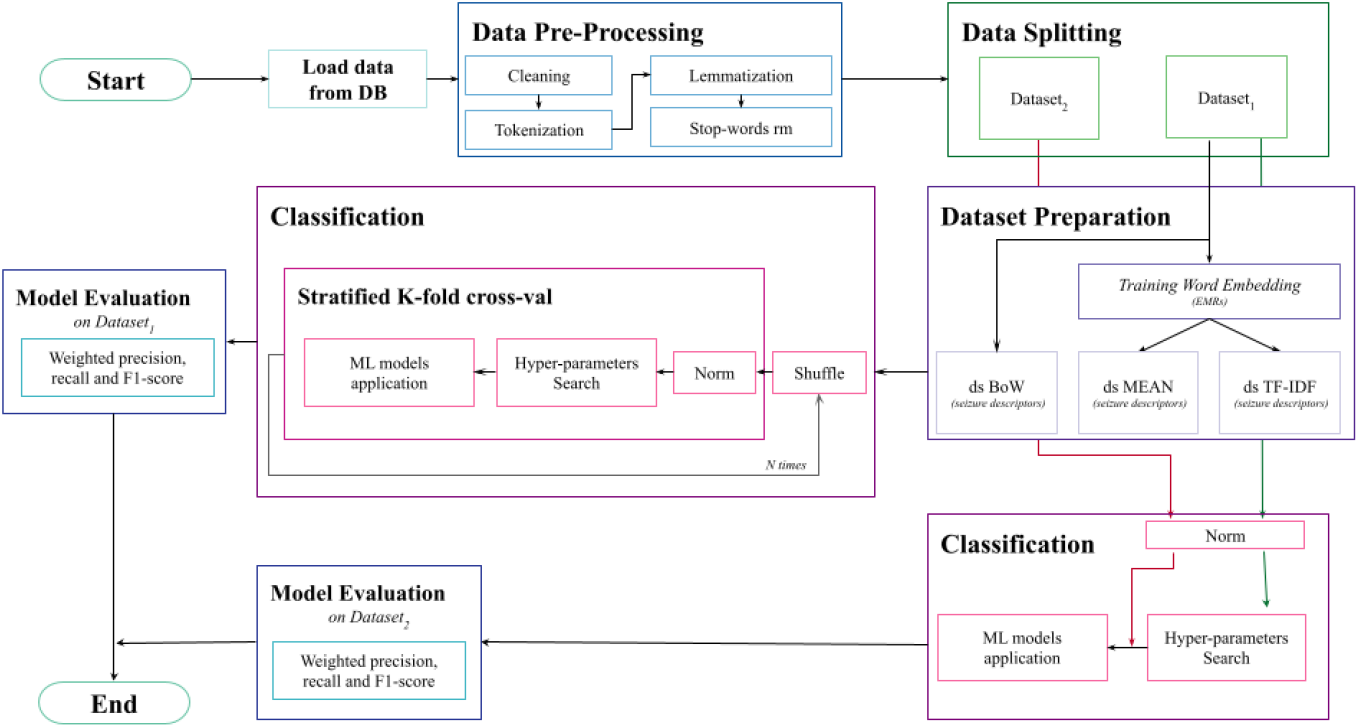
Pipeline Schema. The pipeline can be divided into 5 main sections: data loading, data pre-processing, dataset preparation, classification and model evaluation.

### 2.1 Inclusion criteria

We recruited a large number of subjects with an epilepsy diagnosis from the “Claudio Munari” Epilepsy Surgery Centre, Niguarda Hospital in Milan (Italy). Among them, we selected a group of focal DRE patients who resulted seizure-free after a surgical intervention (minimum follow-up of two years) so that we could exactly identify the origin of seizures (epileptogenic zone). This resulted in a cohort of 129 patients, whose age and sex are distributed according to the pyramid chart in Fig. 2. All participants gave informed consent for data collection and usage for scientific research. This is an anonymous, retrospective, observational study that complies with the principles outlined in the Declaration of Helsinki [24].

**Fig. 2.**
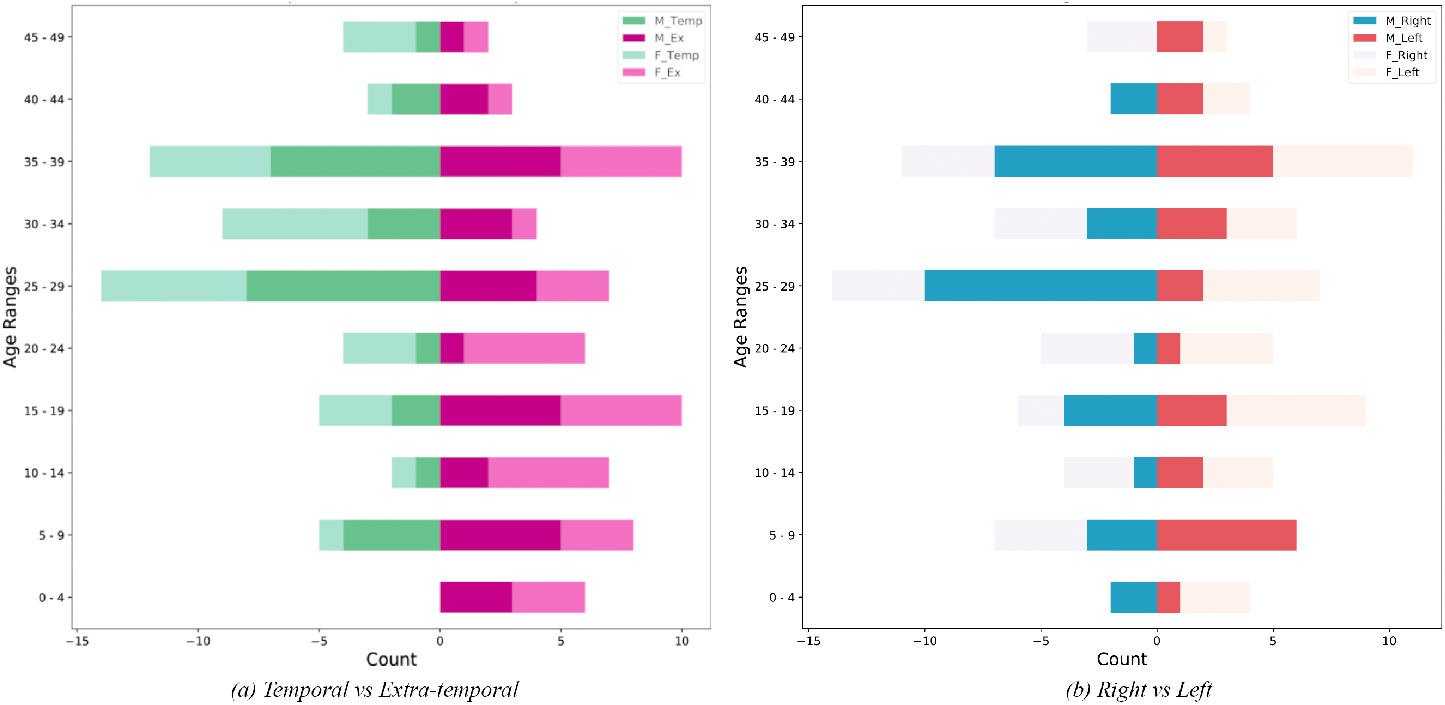
Population distribution. All patients included in the analysis are under 50 years old. In particular, for both tasks temporal/extra-temporal and right/left the majority of the population have an age between 25 and 39.

### 2.2 Samples characteristics

For each patient we collected textual data (consisting of the description of all available seizures, if any, and an excerpt of the patients’ EMR) and the corresponding label that specifies the EZ in the brain. In particular:

#### Seizure descriptions

Texts describing the semiology of seizures. In detail, medical experts watch the recorded videos showing the patient undergoing seizure events and describe the patient’s seizures evolution. Note that one patient may have experienced zero, one or multiple seizures. Specifically, we identified 121 patients with at least one seizure, excluding the descriptions that were labelled as “subjective manifestations” by the experts, for a total of 509 seizure descriptions. The average number of seizures per patient is 4.21±3.51, the maximum is 17. We discarded descriptions that: i) referred to previous seizures (e.g. *“Seizure similar to the previous ones including the automatisms of the right hand brought to the face”*); ii) were composed of less than 20 words; iii) had no mention of physical movements or sensations. In total, we excluded 53 texts resulting in a final dataset of 509 descriptions. We assumed the indipendence of the descriptions for the following reasons: the described seizures occurred in different time spans, the observer who dealt with the patients was not necessarily the same as well as the clinician who wrote the semiological descriptions.

#### EMRs

Anonymous extractions from EMRs from 129 patients, one each. Text files containing several *anamnestic information* such as patient’s history, previous treatments, etc.

The goal of our study is to build predictive models based only on seizure descriptions represented according to some embedding criterion. The problem is cast into a supervised learning setting where each seizure is associated to a label. In our cohort, expert epileptologists identified two types of labels: the region and the side of the brain where the epileptogenic zone is located. This information is available as all patients underwent a surgical intervention that resolved the pathological condition. The first type defines whether the seizure onsets from the temporal (*n*_*temporal*_=58) or extra-temporal (*n*_*extra−temporal*_=63) brain region, where the extra-temporal label also includes patients whose EZ do not cover only the temporal region. The second label type differentiates the EZ based on which hemisphere they are located in (right (*n*_*right*_=61) or left (*n*_*left*_=60)). Considering that, as described above, each patient may suffer from more than one seizure, the dataset is composed of: 39% of seizures labeled in the temporal region and 56% of seizures associated to the right hemisphere.

In order to preserve the morphological structure of the sentences, as well as to anonymize all texts before preprocessing, we removed identificative words, such as given names (substituted with “ragazza/ragazzo” *(“girl”/”boy”)* by matching gender) or cities (substituted with generic “città”*(“city”)*). We stored the anonymized texts in a SQL Server database, using a unique identifier for each patient and retaining only minimal personal information, such as sex and year of birth, as prescribed by the international and national regulations on data protection [25, 26].

### 2.3 Data preparation

For each patient both the EMRs and the available seizure descriptions were stored. To guarantee the semantic interoperability in the classification process of epileptic seizures, e.g. for future studies with more than one center involved, we coded each seizure description with the following main standards used in literature [27]: *International Classification of Diseases 9*^*th*^ *and 10*^*th*^ *edition* (ICD9 and ICD10); ILAE codes [28]. The preprocessing phase followed four main steps:

#### Data cleaning

We cleaned data looking for regular expressions. We removed from the sentences: patterns containing numbers, e.g. dates or names of electrodes, punctuation, text in brackets [29]. Then we extended shared specific abbreviations used by clinicians in their daily practice, e.g. *“aass”* means upper limbs, *“aoo”* means eyes open.

#### Tokenization

We subdivided the content of the text in minimum units of analysis (tokens), which can be single words, or specific groups of words [30]. ***Lemmatization*** We assigned to each word its base form (lemma, e.g. verbs were turned to infinite form and plurals became singular), this way the context of the text was made uniform (normalization process) [31].

#### Stop-words removal

We removed from the text stop-words, i.e. common words (e.g. articles, prepositions, etc.) that are not informative and may alter the building of the models.

### 2.4 NLP & text representation

In order to build a quantitative and meaningful representation of our seizures’ descriptions that can be used as input for learning algorithms, we transformed them from textual data into numerical matrices with two different representation methods:

#### Bag of Words (BW)

A standard sparse representation of text that describes the occurrence of specific n-grams of characters and/or of words within a document. This technique discards any information about the order of words and so the only way to keep the semantic relations between words and to reduce the problem of polysemous words is to use n-grams of words [32]. In this study, we considered all 445 available seizures descriptions in *Dataset*_1_ and we use both n-grams of characters (20%) and n-grams of words (80%) keeping only the most frequent ones within each group. We decided to use n-grams of characters to deal with misspellings, but we left more space to n-grams of words to keep some information about context that otherwise would have been lost. We extracted these n-grams applying the CountVectorizer^2^ method of the Scikit-learn Python library [33] only to the seizure descriptions of the patients belonging to the learning set.

#### Word Embedding

A dense numerical representation of words converted to vectors of numbers that captures the relations between those words based on deep learning. Word embedding can be represented as a function mapping a word *w*, from a vocabulary *V*, to a real-valued vector, in an embedding space of dimension *D*. In our experiments, to generate the embedding, we used the 106 EMRs associated to the 106 patients as well as the 445 Seizure descriptions in *Dataset*_1_. We integrated the information carried by seizure description with the EMRs as they are often composed of syntactically complete sentences and so they are useful to capture the links between words. We used the *Word2vec* model proposed by Mikolov et al. [34] as it has been shown that *Word2vec* better generates word embeddings for most general NLP tasks compared to other approaches [35, 36].

One of the main differences between the two considered approaches is that the Bag of Words model directly returns a representation of the whole document, while the Word Embedding model works at word level. Therefore, when using Word Embedding, we performed a preliminary analysis of the quality of the words’ representation before building the whole document representation. As suggested in [36, 37], we used the following intrinsic evaluators:

*Words similarity* also called *cosine similarity*, which is defined as: norms.

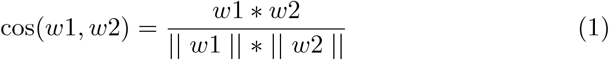

where *w*1 and *w*2 are the two word vectors and || *w*1 || and || *w*2 || are *L*_2_

*Words analogy* given a pair of words (*a* and *a*^*∗*^) and a third word (*b*), the analogy relationship between *a* and *a*^*∗*^ can be used to find the word *b*^*∗*^ that corresponds to *b*.

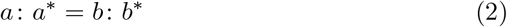

*Outliers detection* given a group of words, the objective is to find the one that does not match the context. This is used to evaluate the semantic coherence in words’ clusters.

Once we obtained the *Word2vec* Word Embedding, we used two approaches to build the representation of seizure descriptions:

*mean representation* In the first one, we averaged all the arrays of the words and we obtained the first matrix.

*tfidf representation* In the second approach, we applied the *Term Frequency-Inverse Document Frequency* (TF-IDF) formula and we obtained the second matrix:

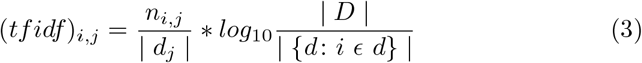

In total we obtained 3 input matrices: the first one using the Bag of Words representation *(bw representation)* and the other two using the Word Embedding model *(mean and tfidf representations)*.

### 2.5 ML methods for classification

For each data representation, we solved a binary classification problem for two different tasks, that are the prediction of the brain region and the prediction of the brain hemisphere where seizures originate. Specifically, we adopted and compared two different machine learning classification methods, that are Logistic Regression with *L*_1_ penalty and SVM, with three different kernels. Hence, in total we had 4 models per each input and task:

#### SVM

A supervised learning method based on maximum margin linear discriminants. The main aim is to find the optimal hyperplane that maximizes the gap or the margin between the classes [13, 14]. To compute the maximum margin search, the loss function that is used in the SVM method is the Hinge Loss [38]. We adopted three different kernels in the SVM classification: *linear, rbf* (gaussian), and *poly* (polynomial with degree equal to 3).

#### Logistic Regression

is a classification technique trained to minimize the Euclidean distance between the model output and the true label. In the specific case of a binary classification, the output variable is modelled by a logistic function (also known as sigmoid) bounded between 0 and 1. An additional *L*_1_ penalty term is used in order to enforce sparsity which allow to build a predictive model based only on a subset of meaningful input variables [15, 16].

### 2.6 Experimental design

#### 2.6.1 Dataset

After the preprocessing phase, the dataset was split into two chunks: *Dataset*_1_ consisting of 445 seizures from 106 patients and *Dataset*_2_ consisting of 64 seizures from 15 patients. The splitting is required to properly assess the generalization properties of the representation/predictive models pairs, as described below.

#### 2.6.2 NLP

Considering the Bag of words representation, we tested three different total number of features: 100, 200, and 300. We obtained best results on the first task considering 200 features while in the second task we obtained best results considering 300 features, from now on we will refer to this numerical representation with *bw*.

Considering the Word Embedding representation, we tested different vector dimensions and combinations of some word embedding parameters for the vector representation, according to reference ranges described in [39]. In particular we ranged: vector dimension between 10, 30, 100; negative sampling between 5, 10, 15; epochs between 100, 200, 300, 400, 500, 600, 700, 800, 900, and 1000. We determined the best combination performing the intrinsic evaluation described in *Sec. NLP & text representations* : vector dimension = 100; negative sampling = 10 and number of epochs = 300. Then we fixed: the minimum words’ occurrence in the text = 2, in order to exclude too rare words or misspellings of frequent words, and number of context words = 3.

#### 2.6.3 ML methods

For all experiments, we performed a stratified k-fold cross-validation, with k = 10, to iteratively split the *Dataset*_1_ into ten different training and validation sets. At each split, the following steps were performed:

1. Data normalization
2. A 10-fold cross-validation to identify the best hyperparameters
3. Model training on the training set with the best hyperparameters
4. Model evaluation on the validation set

All the aforementioned steps have been executed *N* = 3 times, each time shuffling the data. In order to guarantee the reproducibility of the results we set the random state used to shuffle data equal to the iteration index (i.e. in order 0, 1, 2). We then performed an overall evaluation of the model over multiple trials calculating the median performance per trial and the mean performance across the *N* trials.

It is important to specify that we pre-processed seizures of patients belonging to Dataset_2_ together with Dataset_1_ used for cross-validation but we did not use samples from the Dataset_2_ for training the word embedding and the bag of words models. So, when testing generalization ability we looked for the best model hyperparameters using the whole Dataset_1_, while during cross-validation we looked for model hyperparameters only on the training set.

All experiments are evaluated in terms of the following weighted metrics on each fold and on average: precision, recall (which is equal to accuracy), and F1-score. *Sec. Results* describes and graphically represents results in terms of F1-score while the graphs related to the other two metrics are available in the supplementary material.

## 3 Results

### 3.1 Evaluation of the Word Embedding

Mapping high-dimensional data such as natural language text can lead to a spurious representation, so we tested our model with the three intrinsic evaluators described in section “Methods”. We first aimed at verifying if the Word Embedding correctly recognized semantic and syntactic meaning of some random words, so we chose five target words and extracted the most similar words from the Word Embedding according to the Word Similarity measure. We graphically visualized the five target words and the five nearest ones using *T-distributed Stochastic Neighbor Embedding* (T-SNE), which allows the visualization of high dimensional data by locating the datapoints in a two/three-dimensional map [40]. We show that our model correctly associates words with syntactic and semantic meanings similar to the target words in all selected cases *(Fig. 3, Table 1)*. In particular, as we can see in *Fig. 3*, the word “sollevamento *(lift)*” is one of the most similar words to “movimento *(movement)*” but it is also a very similar to “elevazione *(elevation)*” and this property is recognized by the model, as the two words are close together in the two-dimensional projection made by T-SNE. Then, we tested our model with the Words Analogy evaluator, so we wanted to find the word that satisfied the following relation: “braccio *(arm)*” + “gamba *(leg)*” - “piede *(foot)*”. We obtained the expected word “mano *(hand)*”. Finally, we evaluated the ability of outliers detection, so we tested if the Word Embedding model could recognize words out of their context. Specifically, we chose a quadruplet of random words: three within the same context and one outlier. We repeated this experiment three times, finding that our model is always able to detect the outlier *(Table 2)*. Results suggest that our model properly identifies relations between words.

**Fig. 3.**
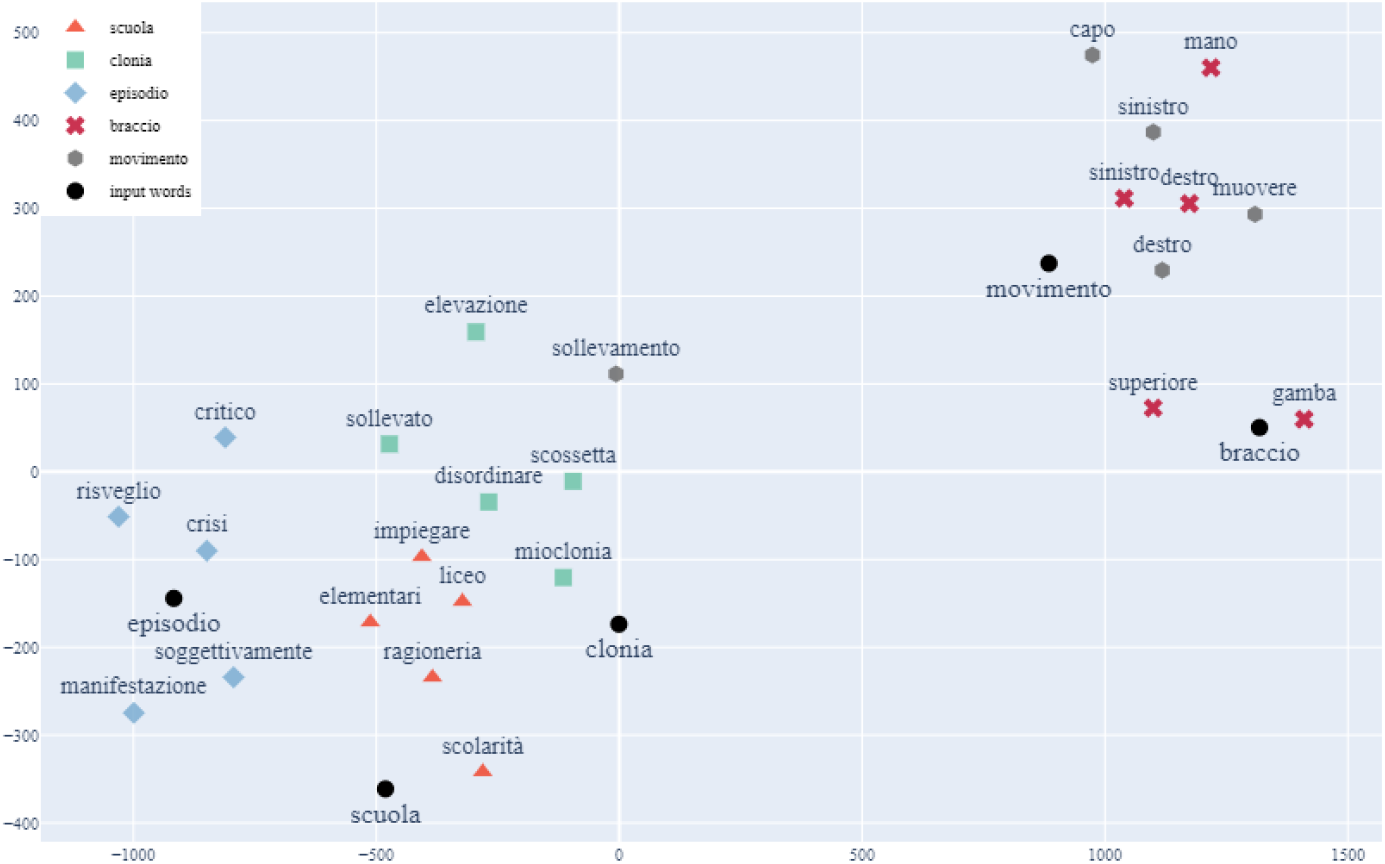
Examples of clusters of Word Embedding, visualized in a two-dimensional space using T-SNE. Each cluster is composed of one input word and the top 5 most similar words. Two main clusters can be distinguished in the figure, words in the upper-right part of the figure are linked to the seizure descriptions (“braccio *(arm)*” and “movimento *(movement)*”) and words in the lower-left part of the figure linked to anamnestic information (“episodio *(episode)*”, “scuola *(school)*” and “clonia *(jerk)*”, which is mentioned most of the times in that section).

**Table 1.** Examples of the words similarity evaluation. The first column contains the five target words arbitrarily selected and the second column contains the corresponding five most similar words induced by the Word Embedding model. Most similar words are written in descending order considering the similarity coefficient based on the cosine similarity.

| Target word | Most similar words |
| --- | --- |
| Braccio<br>(arm) | Gamba, mano, sinistro, destro, superiore<br>(leg, hand, left, right, upper) |
| Episodio<br>(episode) | Crisi, soggettivamente, risveglio, critico, manifestazione<br>(seizure, subjectively, awakening, critical, manifestation) |
| Clonia<br>(jerk) | Mioclonia, scossetta, disordinare, elevazione, sollevato<br>(myoclonia, jerk, clutter, elevation, raised) |
| Scuola<br>(school) | Ragioneria, liceo, elementari, scolarità, impiegare<br>(accounting, high school, primary school, schooling, employ) |
| Movimento<br>(movement) | Muovere, sinistro, destro, capo, sollevamento<br>(move, left, right, head, lift) |

**Table 2.** Examples of the outliers detection evaluator. Given 4 words the trained model could correctly identify the word that did not match the context.

| Input words | Outliers |
| --- | --- |
| Dormire, cuscino, letto, bicchiere<br>(sleep, pillow, bed, glass) | Bicchiere<br>(glass) |
| Scuola, braccio, gamba, mano<br>(school, arm, leg, hand) | Scuola<br>(school) |
| Fratello, sorella, madre, crisi<br>(brother, sister, mother, seizure) | Crisi<br>(seizure) |

### 3.2 Temporal vs. extra-temporal seizure onset sites

The first learning task we faced aimed at defining a predictive model capable of identifying if a given seizure originated into the temporal brain region or elsewhere. In *Figure 4* we report the weighted F1-scores for all combinations of trained model, data shuffle (red, light green, and light blue) and data representation (*mean, tfidf*, and *bw*). Results obtained with Logistic Regression *(Fig. 2a)* show a median F1-score value between 75.26% and 83.95% considering all three representations. The linear SVM model *(Fig. 2b)* showed a median F1-score value between 75.14% and 84.17%. The SVM with rbf kernel *(Fig. 2c)* showed a median F1-score always greater than 75.40% and between 80.96% and 85.42% considering *mean* and *tfidf*. The SVM with poly kernel *(Fig. 2d)* showed a median F1-score value that ranged between 72.72% and 85.27%.

**Fig. 4.**
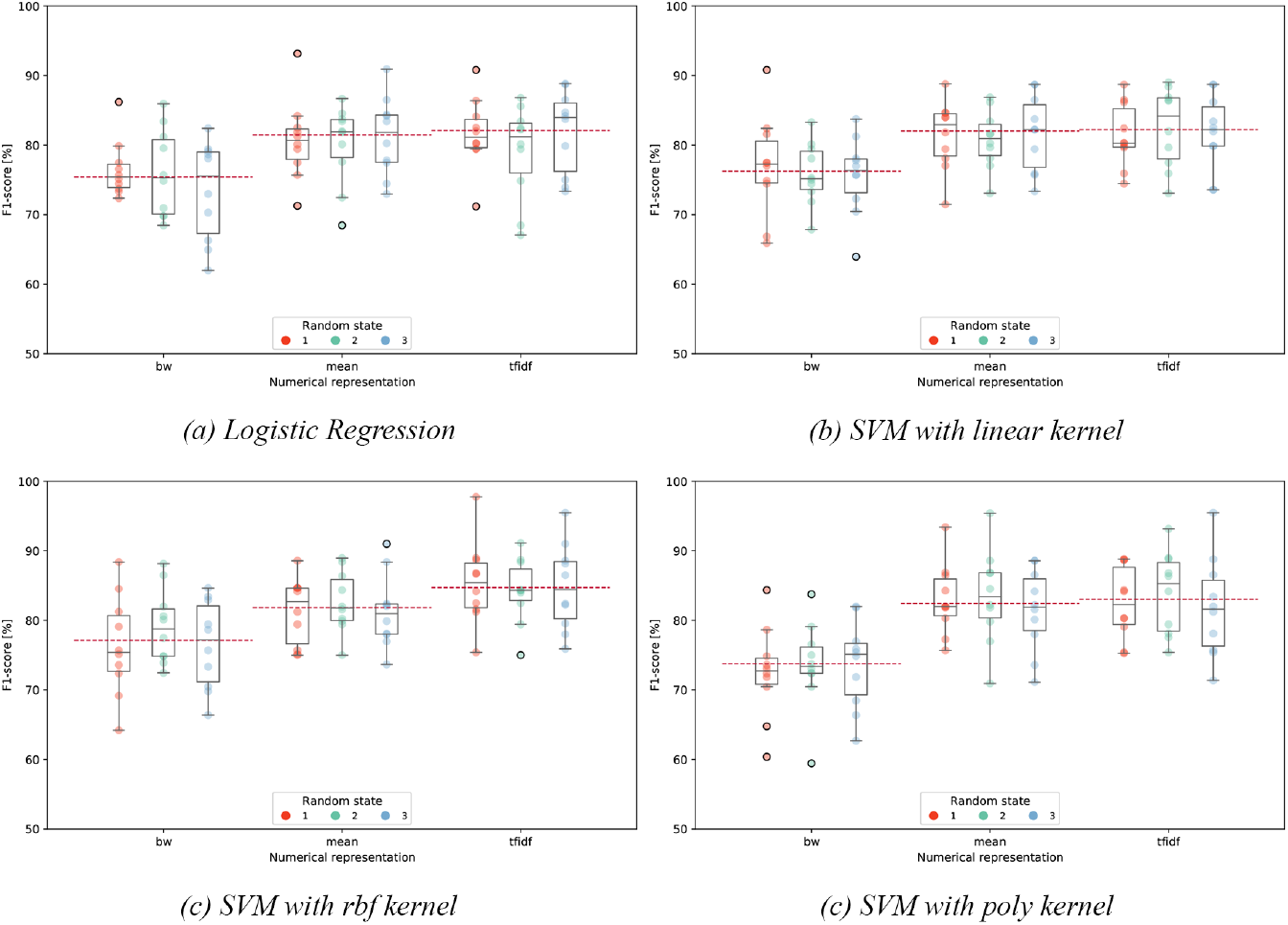
Classification weighted F1-scores in the brain region identification of Logistic Regression (a), SVM with linear kernel (b), SVM with rbf kernel (c), and SVM with poly kernel (d) over the three fixed shuffles and the three numerical representations. For each representation and shuffle the weighted F1-score values of the K-folds are plotted. The three red dotted lines identify the mean of second quartiles over the three shuffles.

All predictive models show a better performance when using the Word Embedding-based representations respect to Bag of Words representation. Between all the combinations of representations and classifiers *(Fig. 5)*, the best performance in terms of F1-score is obtained by the *tfidf* and SVM with rbf kernel.

**Fig. 5.**
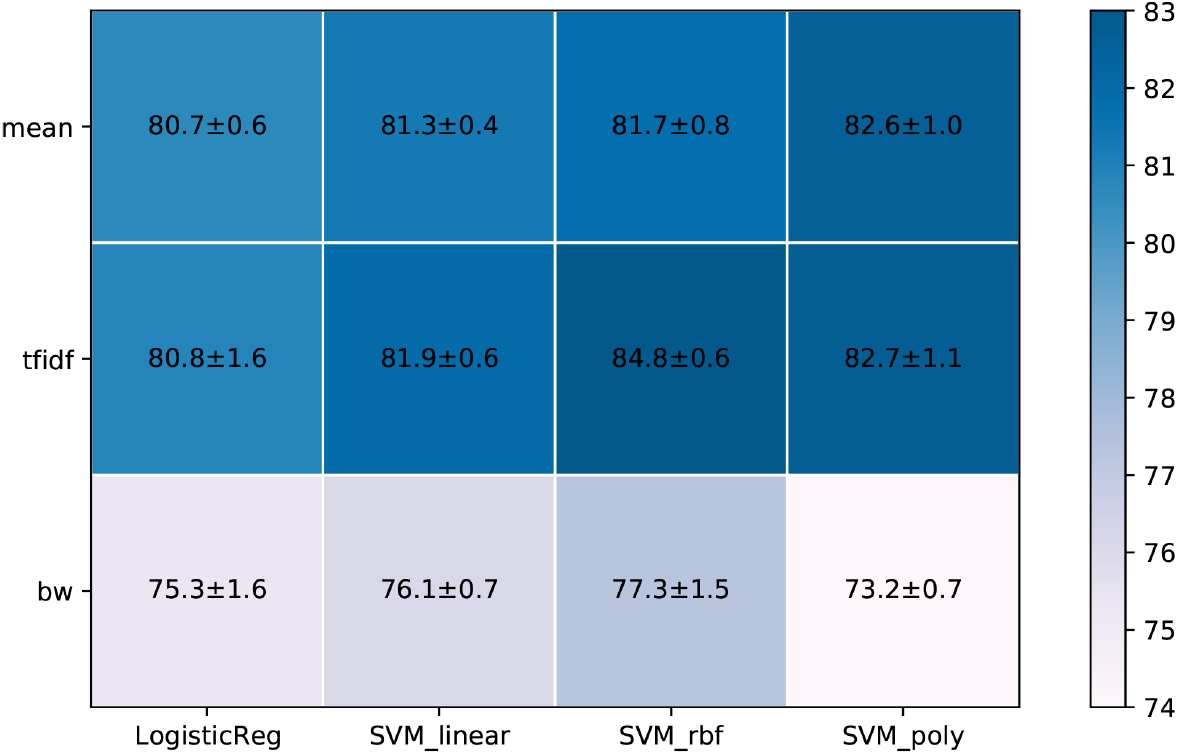
Mean performances in the brain region identification of Logistic Regression, SVM with linear kernel, SVM with rbf kernel, and SVM with polynomial kernel in terms of weighted F1-score. For each numerical representation the mean value of the F1-score is computed from the mean values of each shuffle.

**Fig. 6.**
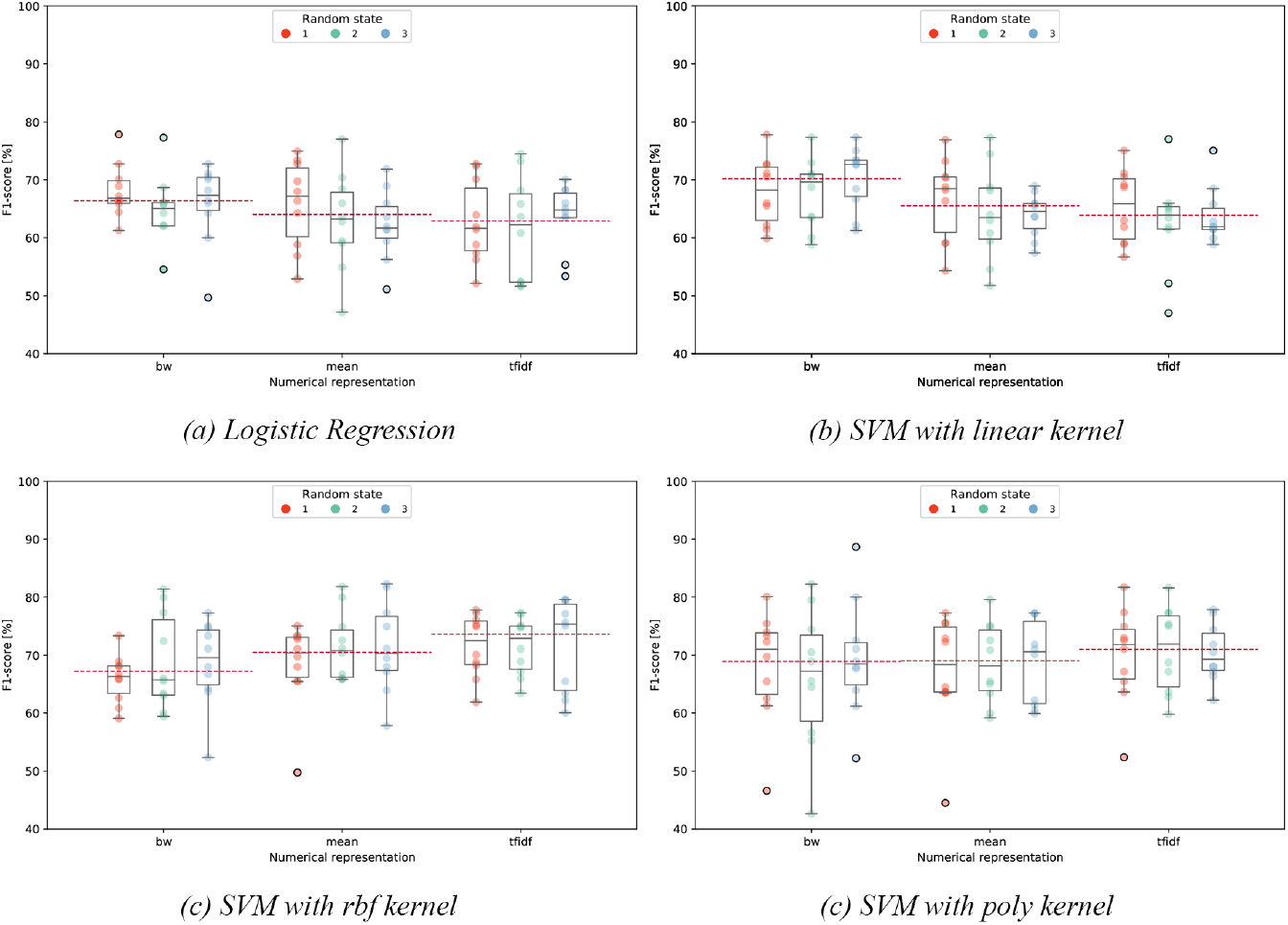
Classification weighted F1-scores on the brain side identification of Logistic Regression (a), SVM with linear kernel (b), SVM with rbf kernel (c), and SVM with poly kernel (d) over the three fixed shuffles and the three numerical representations. For each representation and shuffle the F1-scores of the K-folds are plotted. The three red dotted lines identify the mean of second quartiles over the three shuffles.

**Fig. 7.**
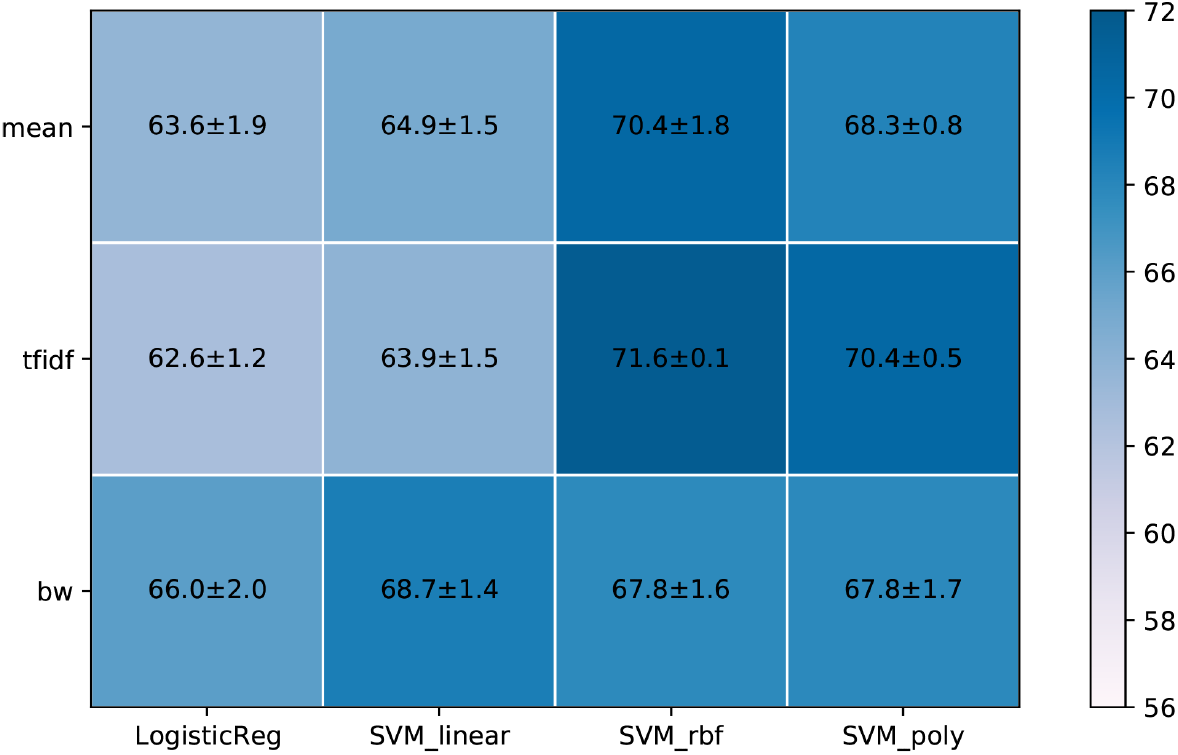
Mean performances in the brain side identification of Logistic Regression, SVM with linear kernel, SVM with rbf kernel, and SVM with polynomial kernel in terms of F1-score over the shuffles and the numerical representations. For each representation the mean value of the F1-score is calculated from the mean values of each shuffle.

We tested our model also on Dataset_2_ in order to assess the ability of generalization of the data representation methods. According to the results reported in *Table 3*, the F1-score on Dataset_2_ obtained with Logistic Regression is 72.13% with *mean*, 74.28% with *tfidf*, and 72.82% with *bw*. Considering the SVM classifier we analyzed the F1-score obtained with each kernel. The linear kernel obtained 73.26% with *mean*, 74.90% with *tfidf*, and 73.07% with *bw*. The rbf kernel showed a value of 78.13% with *mean*, 76.24% with *tfidf*, and it presented 77.72% with *bw*. The poly kernel showed a performance of 70.11% with *mean*, 71.59% with *tfidf*, and 74.54% with *bw*. In general almost all combinations of classification models and numerical representations return a F1-score higher than 70% but the best performances are obtained by SVM with rbf kernel and *mean* and *bw* representations.

**Table 3.** Models performances on Dataset_2_ in the brain region identification. For each combination of numerical representation and classifier the weighted F1-score obtained on the blind test set is reported.

|  | mean | tfidf | BW |
| --- | --- | --- | --- |
| *Logistic regression* | 72.13 | 74.28 | 72.82 |
| *SVM linear* | 73.26 | 74.90 | 73.07 |
| *SVM rbf* | 78.13 | 76.24 | 77.72 |
| *SVM poly* | 70.11 | 71.59 | 74.54 |

### 3.3 Left vs. right hemisphere seizure onset sites

The second learning task aimed at defining a predictive model capable of determining the lateralization of the brain region where seizures originate in the left or right hemisphere. As in the previous task, in *Fig*. we report models weighted F1-scores for all combinations of trained model, data shuffle (red, light green, and light blue) and data representation (*mean, tfidf*, and *bw*). Results obtained with Logistic Regression *(Fig. 4a)* showed a median F1-score value ranging between 61.65% and 67.30% considering all the three representations. The linear SVM model *(Fig. 4b)* showed a median F1-score value ranging between 61.92% and 72.64%. SVM with the rbf kernel *(Fig. 4c)* showed a median F1-score value that is always greater than 65.75% and in particular *mean* and *tfidf* had a median value of F1-score between 70.28% and 75.33%. SVM with poly kernel *(Fig. 4d)* showed a median F1-score that ranged between 67.21% to 71.80%.

## 4 Discussion

Our study shows that a NLP-based tools may help localizing the origin of the seizures in drug-resistant patients candidates to surgery, suggesting a right/left and temporal/extratemporal origin, thus representing a useful tool, especially for those epileptologists without specific skills in the interpretation of seizures-related semeiological data.

A great motivation for the research in this sector derives from the will to speed up the process that leads the patient to be considered a candidate for surgery and also to find alternative types of information that healthcare professionals can collect more easily without necessarily requiring highly specialized diagnostic machinery. Recently, many research projects aimed at developing decision support systems that could provide a more accurate classification of seizures relying on questionnaires [19]. Studies based on self-reported questionnaires are highly dependent on patients’ willingness to personally contribute to the study. Therefore the trend of many research projects moved towards the use of Real World Data, so information already collected and available in electronic format into the EMRs. The main problem with EMRs is that they are mainly composed by unstructured free-text. Hence, the extraction of meaningful information requires a heavy human effort. In order to deal with textual data, many projects introduced NLP techniques and the majority of them focused on information extraction tasks. ExECT [21] is an information extraction system that combines rule-based and statistical techniques to automatically fill the routinely collected data with missing information from epilepsy clinic letters. EpiFinder [20] is a decision support system that extracts keywords to predict the probability to have epilepsy. The pipeline that we presented in this manuscript uses anonymized EMRs and semiological descriptions of seizures, supporting the diagnosis of epilepsy by attempting to ease the process of localization of the epileptogenic zone. In particular, we used NLP methods to pre-process texts and to build numerical representations of the semiological descriptions of individual seizures. We built both count-based and embedding-based models. In particular, we used supervised ML methods to classify the location of seizures origin into right/left hemisphere and temporal/extra-temporal region. Our analysis outlined that, accordingly with literature [35], embedding models obtain best performances on the learning set (*Dataset*_1_). While on unseen data (*Dataset*_2_) good performances are obtained by count-based model as well but this is coherent with the nature of the two numerical representations. The word embedding model is based on deep learning and when applied to *Dataset*_2_ it undergoes an additional process of generalization, compared to when applied to *Dataset*_1_ and so its performances decrease. The main advantage of our pipeline is that we did not adopt specific operation or information linked to the epilepsy context, e.g. we do not rely on ontologies to support models building phase. This renders our pipeline suitable for different clinical scenarios beyond epilepsy. According to our knowledge, our pipeline represents the first NLP-based diagnostic tool for drug-resistant focal epilepsy dedicated to Italian centers. Indeed our project was particularly challenging also because no pre-trained embedding models exist for biomedical applications or other existing works on this topic dealing with Italian language. Our work also presents some limitations. The most important one is that the syntax of EMRs and the semiological descriptions of seizures are highly dependent on the specific clinician writing them. The syntax and choice of terms of each clinician, e.g. the use of different synonyms, impact the building of both the two representations of text. In particular, the training of the word embedding model because each word depends on its context (other near words) and the most relevant features extracted by the count-based model are selected depending on their frequency. Another point to be considered is that we mapped the seizures into national and international coding systems in order to be able in the future to compare the seizures we collected with those coming from other medical centers. Anyway the problem is that terminology standards evolve over time, so in order to guarantee the correct management of the terminology mappings as new versions will be release, we will follow the approach described in [41].

## 5 Conclusions

This work reached encouraging results considering both localization tasks on the testing set of *Dataset*_1_ while only the temporal/extra-temporal task obtained results well above chance level on *Dataset*_2_. However the weak performances of *Dataset*_2_ in the brain side identification task were expected because according to clinicians that task was more difficult compared to the other one. Identifying the EZ represents a challenging task in drug resistant focal epilepsy patients’ assessment. Collectively, these results establish a starting point in the development of a non-invasive, cost-effective tool which can both enhance the pre-surgical evaluation carried out in highly specialized centers and provide a useful support in primary-care units, where many diagnostic procedures may not be available. In both cases this could diminish the delay between epilepsy onset and surgery, with a significant impact on patients’ quality of life and healthcare expenditures.

## List of abbreviations

ASMs: Anti-Seizure Medications
BW: Bag of Words
DRE: Drug-Resistant Epilepsy
EEG: Electro-Encephalogram
EMRs: Electronic Medical Records
ExECT: Extraction of Epilepsy Clinical Text
EZ: Epileptogenic Zone
ICD10: International Classification of Diseases 10th edition
ICD9: International Classification of Diseases 9th edition
ILAE: International League Against Epilepsy
KD: Kawasaki Disease
ML: Machine Learning
MRI: Magnetic Resonance Imaging
NLP: Natural Language Processing
PET: Positron Emission Tomography
SVM: Support Vector Machines
TSNE: T-distributed Stochastic Neighbor Embedding
TFIDF: Term Frequency-Inverse Document Frequency
WHO: World Health Organization

## Declarations

### Ethics approval and consent to participate

All participants gave informed consent for data collection and usage for scientific research. This is an anonymous, retrospective, observational study that complies with the principles outlined in the Declaration of Helsinki [24].

### Consent for publication

Not applicable

### Availability of data and materials

The datasets generated and/or analysed during the current study are not publicly available due to the high personal content of the texts. However the word embedding trained model and the selected list of features of bag of words are available from the corresponding author on reasonable request.

### Author’s contributions

Conceptualization, G.A., A.B., and L.N.; methodology, S.M., R.T., and A.B.; software, S.A. and R.T.; validation, S.M., R.T., and A.B.; formal analysis, S.M.; investigation, S.M.; resources, R.M. and L.T.; data curation, L.C., L.T., R.M., and L.N.; writing—original draft preparation, S.M.; writing—review and editing, S.M., R.T., G.A., A.B., L.C., L.N., L.T., R.M.; visualization, S.M. and R.T.; supervision, G.A. and A.B.; project administration, G.A. All authors read and approved the final manuscript.

## Acknowledgements

DiNOGMI contributed to this work within the framework of the DiNOGMI Department of Excellence MIUR 2018 to 2022 (legge 232/2016).

## Additional Files

**Fig. 8.**
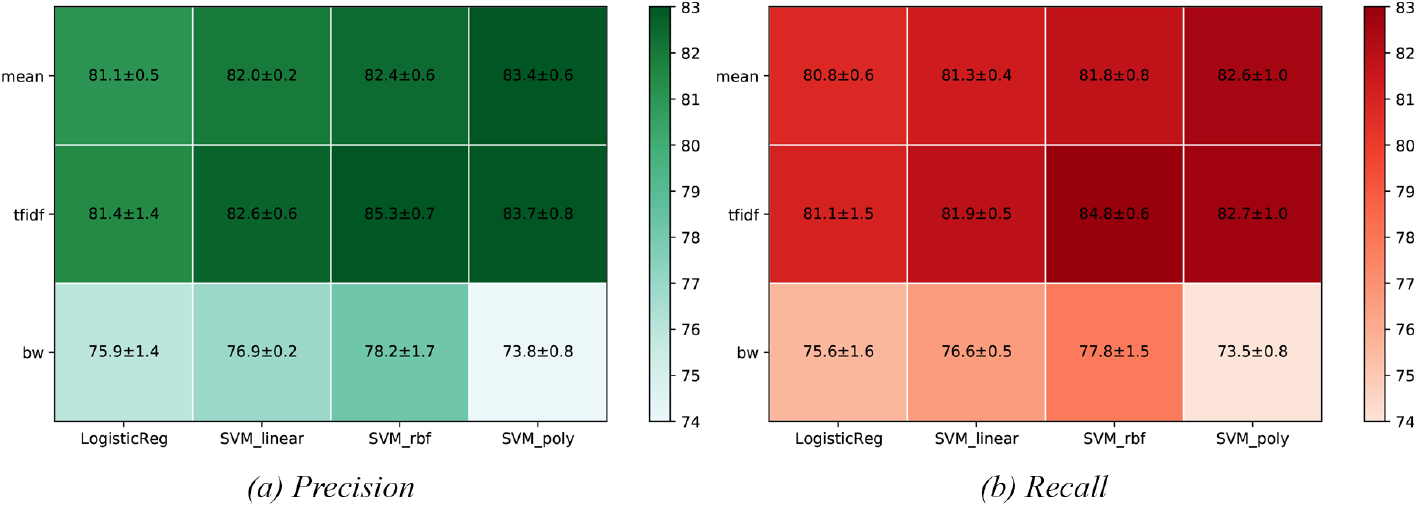
Mean performances in the brain region identification of Logistic Regression, SVM with linear kernel, SVM with rbf kernel and SVM with polynomial kernel in terms of weighted precision and recall over the shuffles and the numerical representations. For each representation the mean value of the specific metric is calculated from the mean values of each shuffle.

**Fig. 9.**
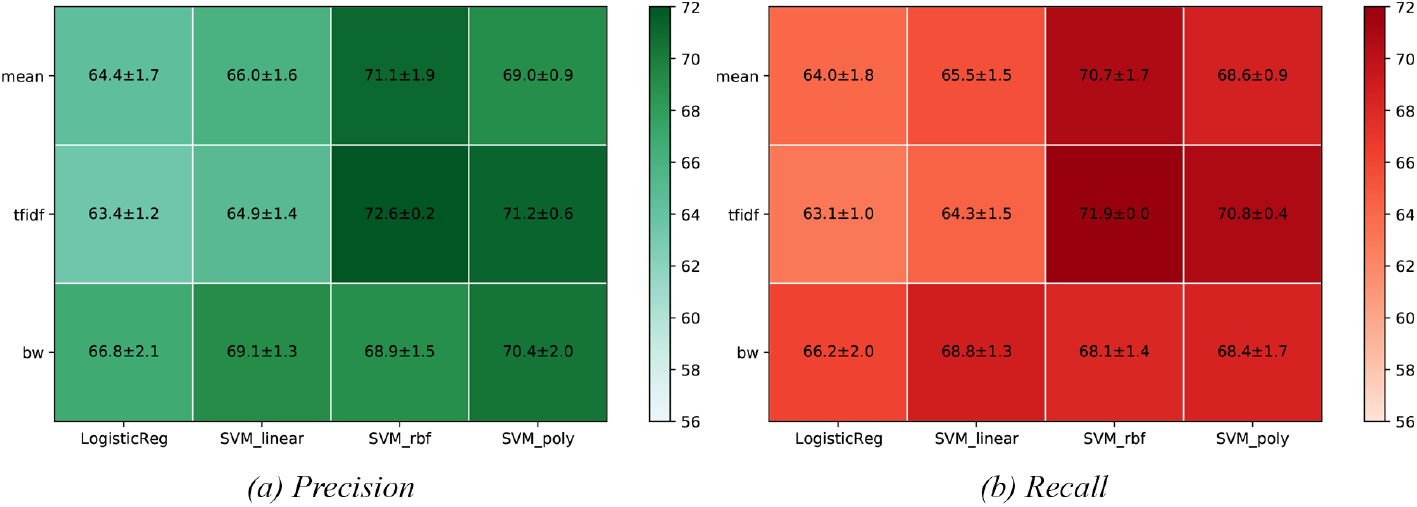
Mean performances in the brain side identification of Logistic Regression, SVM with linear kernel, SVM with rbf kernel and SVM with polynomial kernel in terms of weighted precision and recall over the shuffles and the numerical representations. For each representation the mean value of the specific metric is calculated from the mean values of each shuffle.

## Footnotes

1 https://www.ilae.org/

2 https://scikit-learn.org/stable/modules/generated/sklearn.featureextraction.text.CountVectorizer.html

